# Trajectories of inflammatory biomarkers over the eighth decade and their associations with immune cell counts and epigenetic ageing

**DOI:** 10.1101/397877

**Authors:** Anna J Stevenson, Daniel L McCartney, Sarah E Harris, Adele M Taylor, Paul Redmond, John M Starr, Qian Zhang, Allan F McRae, Naomi R Wray, Tara L Spires-Jones, Barry W McColl, Andrew M McIntosh, Ian J Deary, Riccardo E Marioni

## Abstract

**BACKGROUND:** Epigenetic age acceleration (an older methylation age compared to chronological age) correlates strongly with various age-related morbidities and mortality. Chronic systemic inflammation is thought to be a hallmark of ageing but the relationship between an increased epigenetic age and this likely key phenotype of ageing has not yet been extensively investigated.

**METHODS:** We modelled the trajectories of the inflammatory biomarkers C-reactive protein (CRP; measured using both a high- and low-sensitivity assay), and interleukin-6 (IL-6) over the 8^th^ decade in the Lothian Birth Cohort 1936. We additionally investigated the association between CRP and imputed leukocyte counts. Using linear mixed models we examined the cross-sectional and longitudinal association between the inflammatory biomarkers and two measures of epigenetic age acceleration, derived from the Horvath and Hannum epigenetic clocks.

**RESULTS:** Low-sensitivity CRP declined, high-sensitivity CRP did not change, and IL-6 increased over time. CRP levels inversely associated with total counts of CD8+T cells and CD4+T cells, and positively associated with senescent CD8+T cells, plasmablasts and granulocytes. Cross-sectionally, the Hannum, but not the Horvath, measure of age acceleration was positively associated with low-sensitivity CRP, high-sensitivity CRP, IL-6 and a restricted measure of CRP (≤10mg/L) likely reflecting levels relevant to chronic inflammation.

**CONCLUSIONS:** We found a divergent relationship between inflammation and immune system parameters in older age. We additionally report the Hannum measure of epigenetic age acceleration associated with an elevated inflammatory profile cross-sectionally, but not longitudinally.

## 1. INTRODUCTION

Epidemiological studies have long associated ageing with a progressive move to a chronic inflammatory state, a phenomenon often referred to as ‘inflammaging’ (1, 2). This low-grade, and typically sub-acute, elevation of peripheral pro-inflammatory mediators in the absence of overt infection is strongly associated with the susceptibility to, and progression of, many age-associated diseases such as cancer, type 2 diabetes and Alzheimer’s disease, and is a key risk factor for mortality (3).

Many epidemiological studies of inflammation in older adults have focused on the acute-phase protein C-reactive protein (CRP) and the pro-inflammatory cytokine interleukin 6 (IL-6), both of which are sensitive biomarkers of low-grade inflammation. Akin to other biomarkers, inflammatory mediators can be highly variable yet much of the research into their alterations in ageing has been cross-sectional (4, 5). Multiple-time point measurements are critical to establish the chronicity of inflammatory biomarkers and their trajectories with age, which in itself is key to unravelling the mechanisms behind the process. Several putative pathways have been identified as playing a role in chronic systemic inflammation including increased visceral adiposity, oxidative stress and telomeric and mitochondrial dysfunction (6, 7), and genetic studies have produced candidate polymorphisms associated with the process (8-10). There remains, however, a lack of understanding of the aetiology of chronic inflammation and its relationship with molecular ageing rates (3, 6).

DNA methylation is an epigenetic mechanism by which methyl groups are added to the DNA molecule, typically at Cytosine-Guanine (CpG) dinucleotides. Chronological age has been shown to have a significant effect on methylation levels and, as such, several highly accurate DNA methylation-based biological markers of ageing, or ‘epigenetic clocks’, have been proposed (11-13). Accelerated epigenetic ageing, as evinced by a greater methylation age compared to chronological age, has been linked to various age-associated health outcomes such as frailty (14), lung cancer (15) and Parkinson’s disease (16), as well as all-cause mortality (17, 18). There have been many diverse applications of the epigenetic clock to studies of ageing and disease, but relatively little is known about the relationship between accelerated epigenetic ageing and, a likely key intermediary process of ageing and age-related morbidity: inflammation. Investigating such relationships may give further insight into the molecular aetiology of inflammation and the pathways that regulate it in the ageing process.

In the current study we investigate the dynamics of inflammation across the eighth decade and its relationship with epigenetic age. We established the trajectories of the inflammatory biomarkers CRP and IL-6 and their association with imputed immune cell counts in the Lothian Birth Cohort 1936 (LBC1936). We then assessed their cross-sectional and longitudinal associations with epigenetic age acceleration, using two widely used measures: intrinsic epigenetic age acceleration (IEAA) and extrinsic epigenetic age acceleration (EEAA) (described in methods section). Chronic inflammation is considered to be a pervasive feature of ageing and, as epigenetic age robustly correlates with chronological age, we hypothesise that a faster running epigenetic clock will associate with greater levels of systemic inflammatory biomarkers.

## 2. METHODS

### 2.1 The Lothian Birth Cohort 1936 (LBC1936)

Details of the LBC1936 study have been described previously (19). Briefly, the cohort comprises individuals born in 1936, most of whom who took part in the Scottish Mental Survey 1947, aged about 11 years. 1,091 participants were recruited to the study at a mean age of 70 years and, to date, have completed up to four waves of testing at mean ages of 70, 73, 76 and 79. Data collection for Wave 5 is ongoing (20). At each wave participants have been extensively phenotyped with data obtained on a wide range of health outcomes, lifestyle factors, psycho-social variables, genetics and epigenetics, and cognition.

### 2.2 Ethics and consent

Ethical permission for the LBC1936 was obtained from the Multi-Centre Research Ethics Committee for Scotland (MREC/01/0/56) and the Lothian Research Ethics Committee (Wave 1: LREC/2003/2/29) and the Scotland A Research Ethics Committee (Waves 2, 3 and 4: 07/MRE00/58). Written informed consent was obtained from all participants.

### 2.3 CRP and IL-6

Serum CRP (mg/L) and IL-6 (pg/ml) were measured from venesected whole-blood samples. CRP levels were quantified using both a regular-sensitivity assay (low-sensitivity CRP) at all four waves of data collection, and a high-sensitivity assay (high-sensitivity CRP) at Waves 2 and 3. The low-sensitivity assay was performed using a dry-slide immuno-rate method on OrthoFusion 5.1 F.S analysers (Ortho Clinical Diagnostics). This assay cannot distinguish values less than 3 mg/L and all readings less than 3 mg/L were assigned a value of 1.5 mg/L. The high-sensitivity assay was performed at the University of Glasgow using an enzyme-linked immunosorbent assay (ELISA; R&D Systems, Oxford, UK)(21). IL-6 levels were determined using high-sensitivity ELISA kits (R&D Systems, Oxford, UK) at Waves 2 and 3 (22).

To account for skewed distributions, both CRP and IL-6 levels were log-transformed (natural log) prior to analyses.

### 2.5 Cell counts

Blood cell proportions were estimated based on DNA methylation signatures as described by Chen et al. (17). Briefly, two approaches were used: Houseman’s estimation method and the Horvath method. Houseman’s method uses methylation signatures from purified leukocytes to estimate the abundance of CD8+T cells, CD4+T cells, natural killer cells, B cells and granulocytes (23). The Horvath method, calculated via the advanced analysis option in the above-mentioned epigenetic age calculator software, was used to estimate the percentage of exhausted/senescent CD8+ T cells and plasmablasts (24, 25). Imputed cell counts have been shown to have a moderate correlation with reciprocal flow cytometric data (26).

### 2.4 Epigenetic age acceleration

Methodological details of the methylation profiling for LBC1936 are provided in **Appendix. 1.**

Age acceleration was calculated for each subject at each wave using the online calculator developed by Horvath (https://dnamage.genetics.ucla.edu/)(25).

Two established measures of methylation-based age acceleration were used in this study: intrinsic epigenetic age acceleration (IEAA) and extrinsic epigenetic age acceleration (EEAA). IEAA, based on methylation at 353 CpG sites across multiple tissues as described by Horvath, captures ‘pure’, cell-intrinsic epigenetic ageing, independent of age-related changes in blood cell composition (25). It is defined as the residual resulting from a multivariate regression model of Horvath methylation age on chronological age and estimates of various blood immune cell counts imputed from methylation data.

Conversely, EEAA, an enhanced version of the Hannum clock based on 71CpGs, up-weights the contribution of immune blood cell counts, in addition to tracking intrinsic methylation changes (27). EEAA is calculated through use of a weighted average of Hannum’s methylation age with three cell types that are known to change with age - naïve cytotoxic T-cells, exhausted cytotoxic T-cells and plasmablasts – using the Klerma-Doubal approach (28). EEAA is defined as the residual variation resulting from regressing the weighted estimated age on chronological age (17).

### 2.6 Statistical Analysis

Linear mixed models were used to investigate the baseline relationship between the inflammatory biomarker levels and the imputed cell counts and epigenetic age acceleration measures independently. In each model age, sex and anti-inflammatory drug status (collected at baseline and coded as a dichotomous variable: on medication=1; not on medication=0) were included as covariates. Batch effects (set, position, array, plate and date) were included as random effects to control for technical artefacts.

Mixed effect models were used to examine the longitudinal change in cell counts and inflammatory biomarkers over the four waves for the low-sensitivity CRP, and two waves for the high-sensitivity CRP and IL-6. Here, sex and baseline use of anti-inflammatory medication were included as covariates, age (years) as the timescale. Participant ID and batch effects were fitted as random effects on the intercept. Interaction terms (between chronological age and baseline epigenetic age acceleration, both centred to have mean 0 and SD 1) were included to investigate the changes in inflammatory biomarker levels with age predicted by baseline epigenetic age acceleration. Thus, the model formula was: inflammatory_biomarker ∼ age*baseline_DNAm_age + sex + anti_inflammatory_status + (1|ID) + (1|set) + (1|date) + (1|array) + (1|position) + (1|plate).

Pearson correlations were calculated between IL-6 and the high- and low-sensitivity CRP measures, and between low-sensitivity CRP and the imputed cell counts at each wave separately.

Analyses were performed in RStudio Version 1.0.153 using the ‘lmerTest’ library (29, 30).

## 3. RESULTS

### 3.1 Baseline cohort demographics

Of the 1,091 (543 female; 49.8%) participants recruited at Wave 1, CRP measures (mg/L) were available for 1,054 participants (mean=5.26mg/L, SD=6.68) and epigenetic age acceleration measures for 906 (IEAA: mean=-0.465 years, SD=5.99; EEAA: mean=-0.319 years, SD=7.09). 95 participants were taking anti-inflammatory medication at baseline (8.7%).

### 3.2 Trajectories of inflammatory biomarkers

Spaghetti plots of the trajectories of both raw and log-transformed CRP and IL-6 over time are shown in **Figure 1**. Low-sensitivity log(CRP) was found to decline over the nine years of follow-up by an average of 0.01 standard deviations (SD) per year (p=0.004). The slight increase in high-sensitivity log(CRP) levels over the two waves seen in the plot, was not found to be significant (p=0.718). log(IL-6) increased with age, by an average of 0.15 SD per year (p=2×10^−16^).

**Figure 1.**
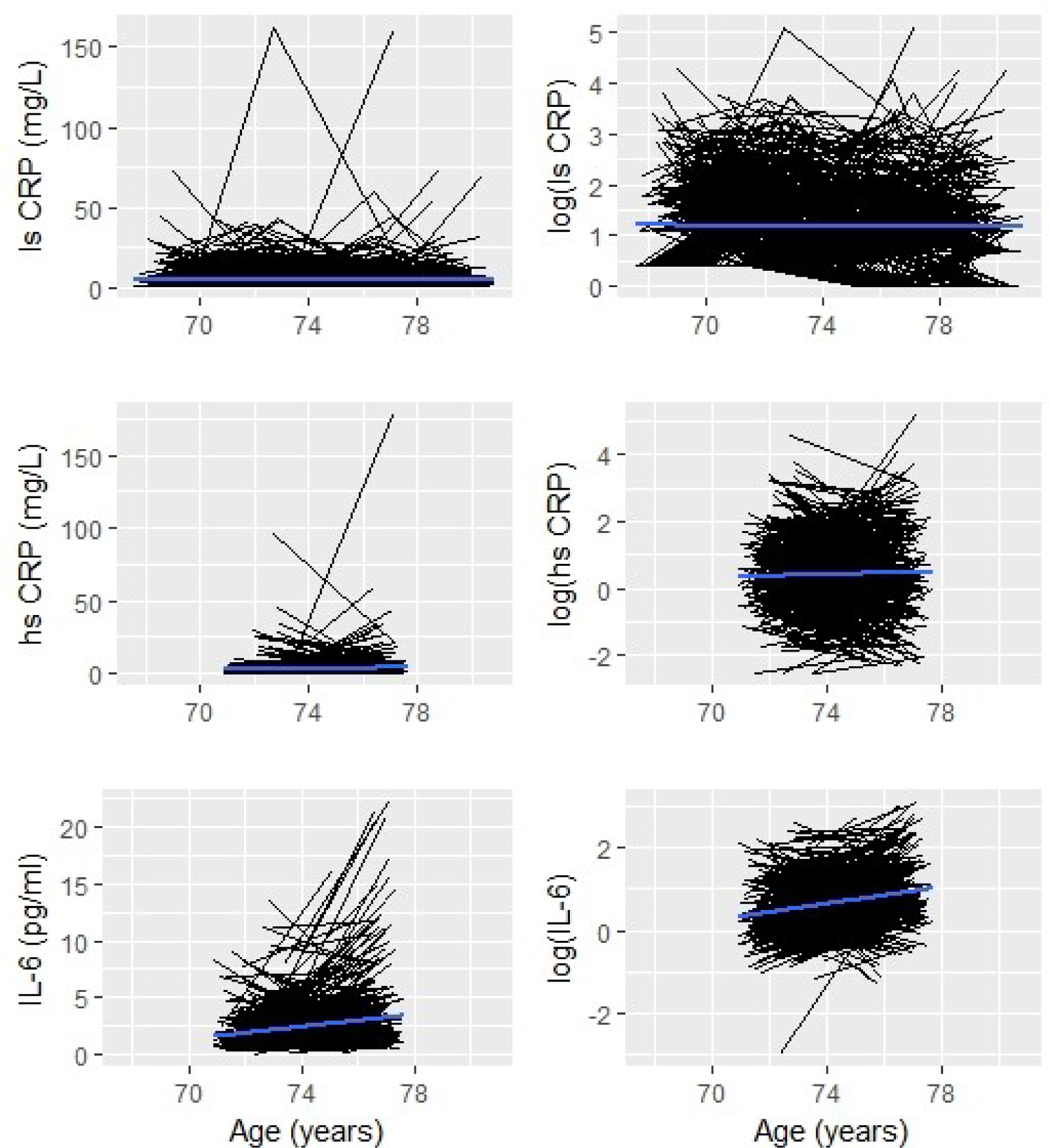
Spaghetti plots of change in CRP and IL-6 over time in LBC1936. CRP: C-reactive protein; IL-6: interleukin-6; hs: high-sensitivity; ls: low-sensitivity; log(): log-transformed.

Due to the discrepancy between the trajectories of the high- and low-sensitivity CRP measures we repeated the analysis of the low-sensitivity measure, restricted to the same time points as were available for the high-sensitivity measure (Waves 2 and 3). Akin to the high-sensitivity measure no change was seen over these two waves (beta=-0.005, p=0.836) indicating that the three year time period between waves is perhaps too brief to capture significant variations in CRP.

### 3.3 Correlations

Correlations between the inflammatory biomarkers both within- and between-waves are presented in **Figure 2**. The high- and low-sensitivity CRP measures correlated strongly within waves (r≥0.93), and both measures had moderate intra-wave correlations with IL-6 (r=0.41-0.45). The low-sensitivity CRP correlations across the four waves were low (r=0.14–0.26). Similar coefficients were seen over the two waves for the high-sensitivity CRP measure (r=0.14) and IL-6 (r=0.33).

**Figure 2.**
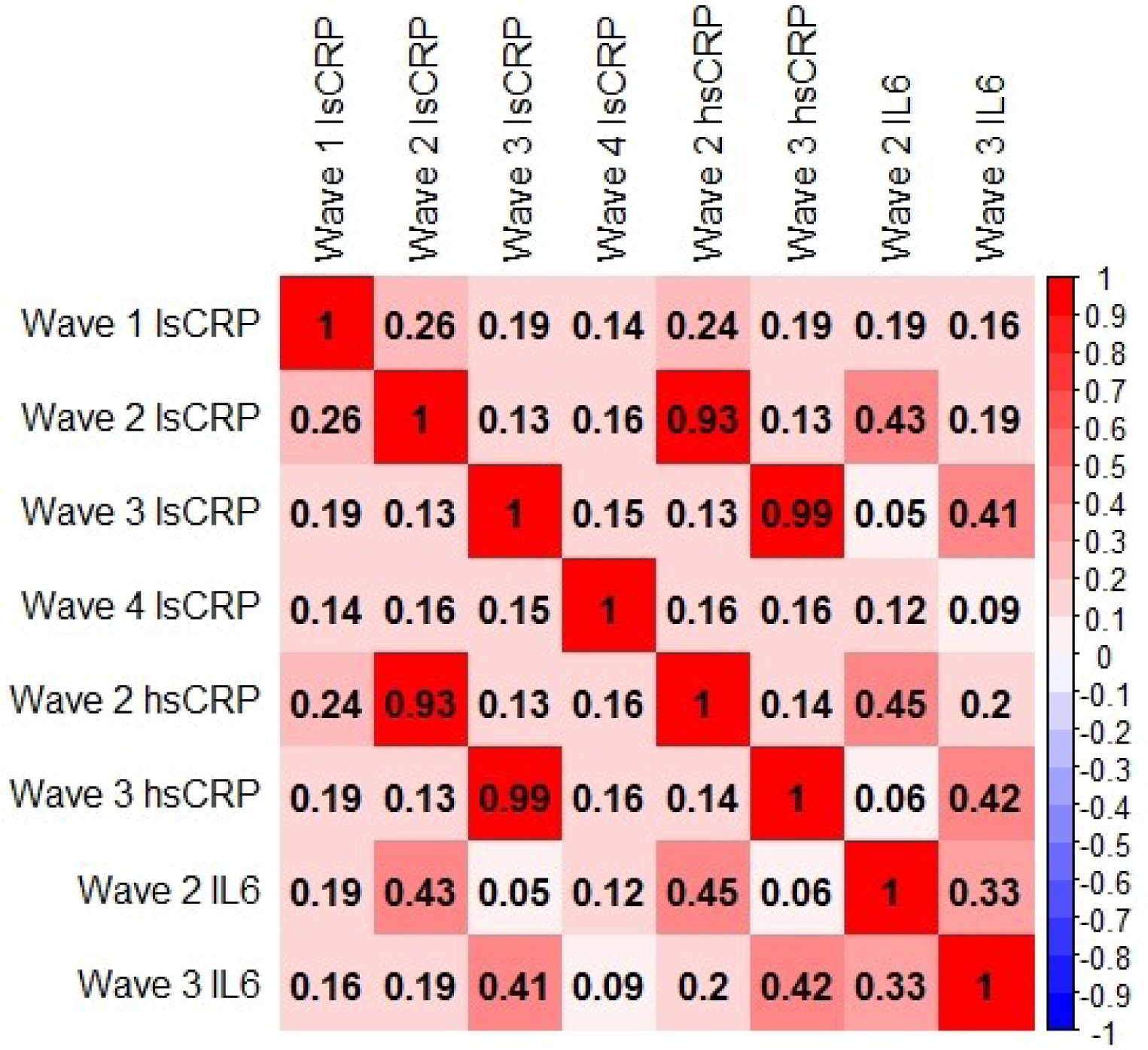
Heatmap of Pearson correlations between inflammatory biomarkers within and across waves of data collection. lsCRP was available at all four waves of data collection; hsCRP and IL-6 were available at Waves 2 and 3 only. CRP: C-reactive protein; IL-6: interleukin-6; ls: low-sensitivity; hs: high-sensitivity.

### 3.5 Associations with imputed cell counts

Due to the association between inflammation and immunosenescence, we examined the dynamics of the immune cell counts imputed from the methylation data, and their association with CRP levels.

We initially established the correlations between both the high- and low-sensitivity CRP measures and the imputed cell counts across the four waves of data (**Table 1**.). There was no significant difference between the correlation coefficients of the two measures across the two waves of overlap (Wilcoxon signed-rank test: wave 2: p=0.76; wave 3: p=0.44) and the directions of the coefficients were consistent. This indicated the lower-sensitivity measure of CRP was representative, and its use in our analyses was unlikely to skew results.

**Table 1.**
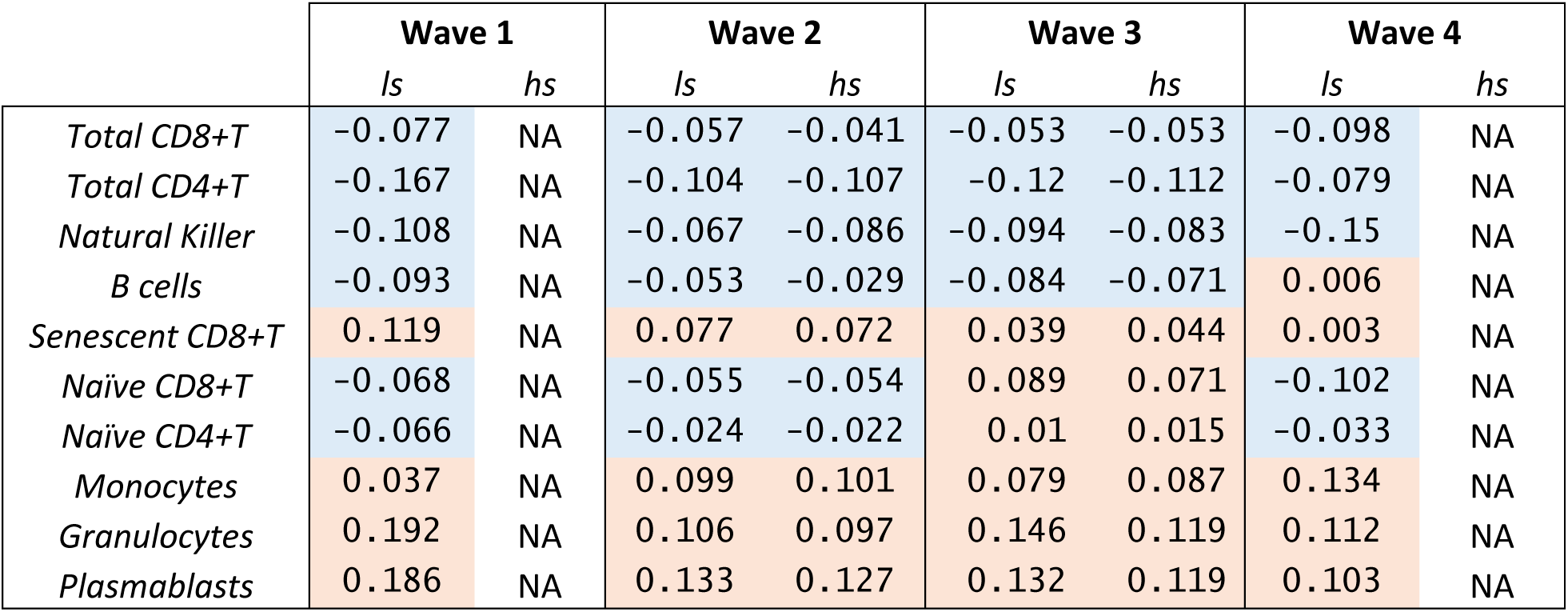
Pearson correlations between high- and low-sensitivity CRP measures and imputed cell counts at all four waves of data. ls: low-sensitivity CRP; hs: high-sensitivity CRP; red: positive correlation; blue: negative correlation.

|  | Wave 1 |  | Wave 2 |  | Wave 3 |  | Wave 4 |  |
| --- | --- | --- | --- | --- | --- | --- | --- | --- |
|  | ls | hs | ls | hs | ls | hs | ls | hs |
| *Total CD8+T* | -0.077 | NA | -0.057 | -0.041 | -0.053 | -0.053 | -0.098 | NA |
| *Total CD4+T* | -0.167 | NA | -0.104 | -0.107 | -0.12 | -0.112 | -0.079 | NA |
| *Natural Killer* | -0.108 | NA | -0.067 | -0.086 | -0.094 | -0.083 | -0.15 | NA |
| *B cells* | -0.093 | NA | -0.053 | -0.029 | -0.084 | -0.071 | 0.006 | NA |
| *Senescent CD8+T* | 0.119 | NA | 0.077 | 0.072 | 0.039 | 0.044 | 0.003 | NA |
| *Naïve CD8+T* | -0.068 | NA | -0.055 | -0.054 | 0.089 | 0.071 | -0.102 | NA |
| *Naïve CD4+T* | -0.066 | NA | -0.024 | -0.022 | 0.01 | 0.015 | -0.033 | NA |
| *Monocytes* | 0.037 | NA | 0.099 | 0.101 | 0.079 | 0.087 | 0.134 | NA |
| *Granulocytes* | 0.192 | NA | 0.106 | 0.097 | 0.146 | 0.119 | 0.112 | NA |
| *Plasmablasts* | 0.186 | NA | 0.133 | 0.127 | 0.132 | 0.119 | 0.103 | NA |

All lymphocyte cell counts declined over time: CD8+T total (beta=-0.01 SD/year, p=4.46 × 10^−7^); naïve CD8+T (beta=-0.01 SD/year, p=4.22 × 10^−5^); total CD4+T (beta=-0.06 SD/year, p=2 × 10^−16^); naïve CD4+T (beta=-0.05 SD/year, p=2 × 10^−16^), NK (beta=-0.04 SD/year, p=2 × 10^−16^); B (beta=-0.04 SD/year, p=2 × 10^−16^). Exhausted/senescent CD8+T cells increased with age (beta=0.03 SD/year, p=1.18 × 10^−14^) as did monocytes (beta=0.06 SD/year, p=2×10^−16^), granulocytes (beta=0.04 SD/year, p=2 × 10^−16^) and plasmablasts (beta=0.08 SD/year, p=2 × 10^−16^ **Figure 3**.)

**Figure 3.**
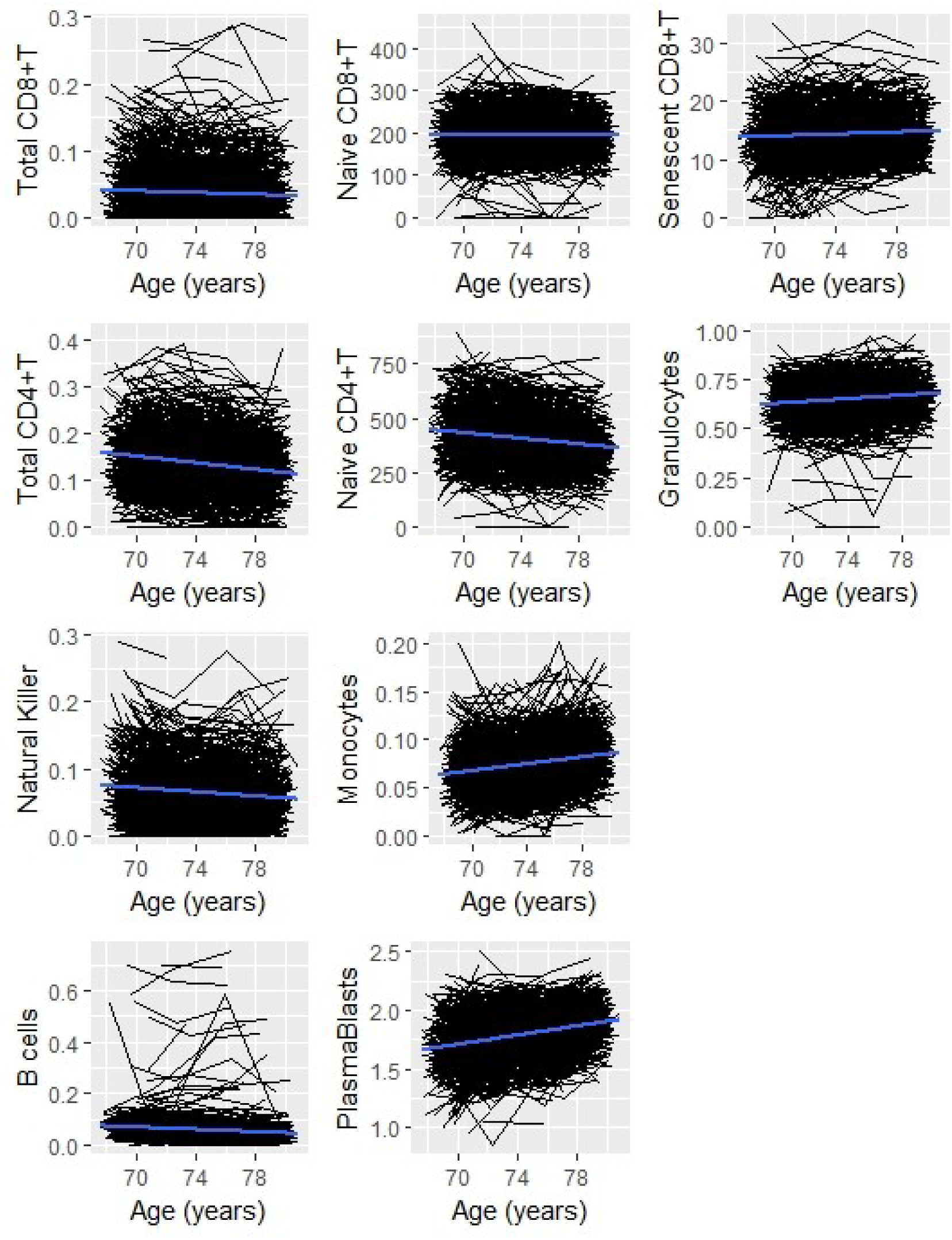
Spaghetti plots of change in imputed immune cell counts over time.

The baseline associations between the blood cellular composition and low-sensitivity log(CRP) within the cohort are presented in **Table 2**. An inverse association was observed between CRP and CD8+T cells (total: beta=-0.073, p=0.026; naïve: beta=-0.142, p=2.54 × 10^−5^) and CD4+T cells (total: beta=- 0.132, p=6.74 × 10^−5^, naïve: beta=-0.072, p=0.032). Higher levels of CRP were associated with greater senescent CD8+T cells (beta=0.134, p=3.79 × 10^−5^), plasmablasts (beta=0.148, p=1.38 × 10^−5^) and granulocyte counts (beta=0.133, p=4.17 × 10^−5^). No associations were found between log(CRP) and B cells, NK cells, or monocytes. We ran a sensitivity analysis excluding those with CRP>10mg/L to test if the associations were altered with more chronic levels of inflammation. Here, the associations with CD4+T cells, total CD8+T cells and granulocytes were attenuated, though the coefficients were in the same direction as previously. All other results remained the same though with larger p-values (**supplementary Table 1**.).

**Table 2.** Baseline associations between low-sensitivity C-reactive protein and imputed cell counts. Significant associations are highlighted in bold.

|  | *Standardised* |  |  |
| --- | --- | --- | --- |
|  | *beta* | *SE* | *P* |
| *Total CD8+T* | -0.073 | 0.033 | **0.026** |
| *Total CD4+T* | -0.132 | 0.033 | **6.74 x 10<sup>-5</sup>** |
| *Naïve CD8+T* | -0.142 | 0.033 | **2.54 x 10<sup>-5</sup>** |
| *Naïve CD4+T* | -0.072 | 0.034 | **0.032** |
| *B cell* | -0.063 | 0.033 | 0.052 |
| *Natural Killer* | -0.055 | 0.033 | 0.094 |
| *Senescent CD8+T* | 0.134 | 0.033 | **3.79 x 10<sup>-5</sup>** |
| *Monocytes* | 0.052 | 0.034 | 0.123 |
| *Plasmablasts* | 0.148 | 0.034 | **1.38 x 10<sup>-5</sup>** |
| *Granulocytes* | 0.133 | 0.033 | **4.17 x 10<sup>-5</sup>** |

### 3.4 Associations with age acceleration

We assessed the cross-sectional and longitudinal relationship between each inflammatory biomarker and methylation-based age acceleration using IEAA and EEAA (**Table 3**). Positive baseline associations were found between both the low- and high-sensitivity CRP measures and EEAA (low-sensitivity log(CRP): beta=0.154, p=2.48 × 10^−5^, high-sensitivity log(CRP): beta=0.111, p=0.004). No baseline correlations were seen between either CRP measure and IEAA (p≥0.087). Neither baseline IEAA nor EEAA was associated with longitudinal change in log(CRP) (p≥0.360).

**Table 3.** Baseline (Wave 1 for low-sensitivity CRP; Wave 2 for high-sensitivity CRP and IL-6) and longitudinal associations between log-transformed inflammatory biomarkers and epigenetic age acceleration. Significant associations are highlighted in bold. CRP: C-reactive protein; IL-6: interleukin 6.

| Baseline | IEAA* |  |  | EEAA† |  |  |
| --- | --- | --- | --- | --- | --- | --- |
|  | Standardised<br>beta | SE | p | Standardised<br>beta | SE | p |
| Low-sensitivity CRP | 0.057 | 0.033 | 0.087 | 0.154 | 0.034 | **2.48x10<sup>-5</sup>** |
| High-sensitivity CRP | 0.047 | 0.036 | 0.189 | 0.111 | 0.037 | **0.004** |
| IL-6 | 0.021 | 0.036 | 0.563 | 0.114 | 0.037 | **0.003** |
| Low-sensitivity CRP ≤10mg/L | 0.062 | 0.035 | 0.081 | 0.129 | 0.039 | **0.0009** |
| **Longitudinal** |  |  |  |  |  |  |
| Low-sensitivity CRP | -0.017 | 0.019 | 0.360 | -0.008 | 0.019 | 0.672 |
| High-sensitivity CRP | 0.005 | 0.039 | 0.888 | -0.006 | 0.039 | 0.888 |
| IL-6 | 0.034 | 0.041 | 0.392 | 0.009 | 0.042 | 0.829 |
| Low sensitivity CRP ≤10mg/L | -0.020 | 0.020 | 0.303 | -0.006 | 0.020 | 0.759 |

\* intrinsic epigenetic age acceleration
† extrinsic epigenetic age acceleration

log(IL-6) also showed a positive baseline association with EEAA (beta=0.114, p=0.003) but not with IEAA (p=0.563). There was no longitudinal association between the log(IL-6) measure and either measure of epigenetic age acceleration (p≥0.392).

A CRP level of ≤10mg/L is often cited as reflecting the non-acute, chronic inflammation, relevant to what has been reported in ageing (31). Because of this, we additionally investigated the relationship between epigenetic age acceleration and log(CRP) in the subset of the cohort with CRP levels of ≤10mg/L, as measured by the low-sensitivity assay (**Table 3**). A positive cross-sectional association was present between EEAA and CRP levels (beta=0.129, p=0.0009). No equivalent association was observed with IEAA (p=0.081). There was no longitudinal associations with either measure of epigenetic age acceleration (p≥0.303).

## 4. DISCUSSION

We found a decline over time in the serum levels of the inflammatory mediator CRP, measured using a low-sensitivity assay in LBC1936. The CRP levels measured from the high-sensitivity assay did not exhibit similarly significant changes with age, whereas the pro-inflammatory cytokine IL-6 was found to increase in the cohort across the two waves of follow up from age 73 to 76. We found a decline over time in naïve and total CD8+T cells, naïve CD4+T cells, NK cells and B cells. Congruent with flow cytometric data, senescent CD8+T cells were found to increase with age as did monocytes, granulocytes and plasmablasts [17, 38, 39]. Negative baseline associations were found between CRP and both naïve and total CD8+T cells and CD4+T cells, and positive associations between CRP and senescent CD8+T cells, plasmablasts and granulocytes. We report associations between both high- and low-sensitivity measures of CRP, IL-6 and a restricted CRP measure (≤10mg/L) with extrinsic estimates of epigenetic age acceleration. These findings suggest a higher epigenetic age is associated with an increased inflammatory profile, in relation to both general inflammation, and to levels likely reflecting chronic, low-grade inflammation.

The dynamics of the low-sensitivity CRP in the cohort were paradoxical to what we might have predicted in a longitudinal study of ageing as previous studies have shown a progressive increase in CRP levels as a function of age (7, 32). Furthermore, IL-6 is a major driving factor in CRP production yet, despite observing a rise in the cytokine over Waves 2 and 3, no corresponding increase is seen in the low-sensitivity CRP levels; in fact the opposite is seen. These findings may be due to the poor sensitivity of the assay used to quantify the CRP levels, particularly at the lower range of values, though levels were found to correlate strongly with those from the more sensitive measure. It is possible that a disruption in the pathway of CRP production impacted our results. CRP is of hepatic origin and it is feasible that impaired liver function, which is common in older adults, influenced its production within individuals in the cohort. Similarly, CRP production is influenced by other cytokines alongside IL-6, such as IL-1 and IL-17, and as these were not measured in the current study their potential influence is unknown (33). Another explanation may lie in the fact the LBC1936 is a largely healthy older ageing cohort. Multiple chronic conditions are common in elderly populations and our results may reflect a more successful ageing process revealed through more static or declining CRP levels. Though part of the innate immune system, inflammation may be a somewhat adaptive response in older age and research has indicated it might confer protection when under tight control (31, 34). Indeed, centenarians often have signs of systemic inflammation but are not afflicted by age-associated diseases (35). It is often argued that inflammation is beneficial in neutralising harmful stimuli in early life but becomes detrimental in older age; however, it may be that optimum levels of inflammation change, but continue to exist, in old age and excessive, insufficient, or highly fluctuating, levels could exacerbate the risk of disease. Finally, the mean age of the cohort at Wave 1 was 70 years and it is possible participants had already undergone a transition to an increased inflammatory profile and this is why we did not capture a rise in CRP over time. This is in line with findings from a previous study which reported an increased serum level of CRP between a 20-30-year-old age group and 60-70-year-olds, but no significant difference in levels between 60-70-year-olds and the 70-90-year-olds (36).

Because of the complexity in the inflammatory trajectories, and given that adaptive immune dysregulation is thought to be an important reciprocal mechanism to age-related inflammation, we additionally examined the dynamics of the immune cell counts imputed from the methylation data, and their association with CRP. The baseline associations between CRP and the cell counts were not congruent, indicating the relationship between inflammatory mediators and individual immune cells is not uniform, with different compartments of the system displaying different relative characteristics. The interrelationship between inflammation and the immune system is clearly complex and the dyad is probably influenced further by other additional pathophysiological pathways activated in older age. Our results fit with the remodelling theory of ageing which postulates that immunosenescence or ‘immuno-remodelling’ is a dynamic process involving both loss and gains of immune function (34, 37). This theory hypothesises that those with a superior capacity to adapt their inflammatory and immune responses, rather than necessarily generating the most robust response, age most successfully. Exactly which of these alterations may be beneficial, and which detrimental, remains to be determined.

Despite no evidence of an increase in CRP with chronological age within the cohort, we found positive associations between all inflammatory biomarkers and extrinsic epigenetic age acceleration, suggesting a faster running epigenetic clock is associated with an increased inflammatory profile. No parallel association was seen between any of the inflammatory mediators and the intrinsic (cell-adjusted) measure of age acceleration. This discordance may be due to the difference in the two estimates. IEAA is calculated independently of the blood cellular composition, measures cell-intrinsic methylation changes, and likely captures an ageing process that is mostly conserved across cell-types. Contrastingly, EEAA does capture age-related alterations in leukocyte composition and correlates with health-related characteristics (38). Evidently, systemic inflammation is closely tied to the blood tissue and so is perhaps more likely to be discerned by a blood-based measure than one that focuses on multiple tissue types. Our results are similar to a recent cross-sectional study on a cohort of middle-aged adults where EEAA was found to correlate strongly with CRP and IL-6 but no similar association was found with IEAA (39).

The main strength of this study is its basis in a relatively large sample of older people with long-term follow-up. The repeated measures of the serum biomarkers has permitted both a description of their trajectories across the eighth decade, and the investigation of longitudinal associations with biological age. The study may have been limited by the cohort effect; as discussed above, the sample is representative of healthier older ageing and findings may not be relevant to typical ageing in which multi-morbidity is common. Additionally, we only examined a restricted number of inflammatory variables and the observed associations with epigenetic age acceleration do not provide evidence of causation. As the trajectories of the inflammatory biomarkers were somewhat complex it is difficult to determine which is most representative. IL-6, which showed a significant increase, may be a more indicative measure of inflammation, but as it was only available for two time-points we focused our analyses on CRP. Recent results have questioned the utility of DNA methylation patterns as biomarkers of aging, as a reduced prediction error of chronological age was found in studies using larger sample sizes to train the age predictor (40). These findings indicate a limitation in the variation in biological age that is captured by DNA methylation and caution is needed in the interpretation of results from epigenetic age predictors. Finally, we did not apply a correction for multiple testing. Applying a strict Bonferonni-corrected threshold of significance would result in the attenuation of the less highly-significant finding of the association between total CD8+T cells and CRP.

In conclusion, the dynamics of the assessed inflammatory markers did not conclusively confirm an increased inflammatory state with older chronological age. We found, however, that a faster running epigenetic clock, as measured by extrinsic age acceleration, associated with a raised inflammatory profile. This association should be investigated further with respect to causal inference. The relationship between CRP and imputed immune cell counts suggest a divergent association between inflammatory mediators and immune parameters. Whether these are adaptive responses or detrimental changes is unclear should be addressed by future studies.

## ACKNOWLEDGEMENTS

The authors thank all LBC1936 study participants and research team members who have contributed, and continue to contribute, to the ongoing LBC1936 study. LBC1936 is supported by Age UK (Disconnected Mind programme) and the Medical Research Council [MR/M01311/1]. Methylation typing was supported by the Centre for Cognitive Ageing and Cognitive Epidemiology (Pilot Fund award), Age UK, The Wellcome Trust Institutional Strategic Support Fund, The University of Edinburgh, and The University of Queensland.

